# Biochemical, Biophysical, and Functional Analyses of Two Isoforms of the SnRK2 inhibitor AtSCS

**DOI:** 10.1101/813717

**Authors:** Krzysztof Tarnowski, Maria Klimecka, Arkadiusz Ciesielski, Grażyna Goch, Anna Kulik, Halina Fedak, Jarosław Poznański, Małgorzata Lichocka, Marcin Pierechod, Richard A. Engh, Michał Dadlez, Grażyna Dobrowolska, Maria Bucholc

## Abstract

SNF1-related protein kinases 2 (SnRK2s) are key signaling elements that regulate abscisic acid (ABA)-dependent plant development and responses to environmental stresses. Our previous data showed that the SnRK2-interacting Calcium Sensor (SCS) is an inhibitor of SnRK2 activity. In *Arabidopsis thaliana*, the use of alternative transcription start sites located within *AtSCS* gene results in two in-frame transcripts and subsequently two proteins, which differ only by the sequence position of the N-terminus. We described the longer AtSCS-A earlier, and now we describe the shorter AtSCS-B and compare both isoforms. The two forms differ significantly in their expression profiles in plant organs and in response to environmental stresses, in calcium binding properties, and conformational dynamics in the presence and absence of Ca^2+^. The results show that only AtSCS-A has the features of a calcium sensor. Both forms inhibit SnRK2 activity, but differ with respect to calcium dependence, as AtSCS-A requires calcium for inhibition, while AtSCS-B does not. Analysis of Arabidopsis plants stably expressing *35S::AtSCS-A-c-myc* or *35S::AtSCS-B-c-myc* in the *scs-1* knockout mutant revealed that *in planta* both forms are negative regulators of the SnRK2 activity induced in response to ABA and regulate plant defense against water deficit. Moreover, the data present biochemical, biophysical, and functional properties of EF-hand-like motifs in plant proteins.

**One sentence Summary:** Two isoforms of SnRK2-interacting calcium sensor are expressed in Arabidopsis; they differ in calcium binding properties, but both of them inhibit SnRK2s and subsequently fine tune ABA signaling.

## INTRODUCTION

SNF1-related protein kinases 2 (SnRK2s) are plant specific kinases involved in plant response to environmental stresses (e.g., water deficit, salinity) and in abscisic acid (ABA)-dependent development (for reviews see: Hubbard et al., 2010; Fujita et al., 2011; Kulik et al., 2011; Nakashima and Yamaguchi-Shinozaki, 2013; Yoshida et al., 2015). Based on phylogenetic analyses, SnRK2s have been classified into three groups. The classification correlates closely with their sensitivity to ABA; group 1 consists of SnRK2s which are not activated by ABA treatment, group 2 includes kinases which are not or only weakly activated by ABA, and group 3 kinases are strongly activated by the phytohormone. Ample data demonstrate the role of group 3 SnRK2s in ABA signaling, both in plant development as well as in stress response. In *Arabidopsis thaliana*, group 3 comprises three members: SnRK2.2, SnRK2.3 and SnRK2.6. SnRK2.2 and SnRK2.3 are involved mainly in the regulation of seed dormancy and germination (Fujii *et al*., 2007), whereas SnRK2.6 regulates stomatal closure in response to water deficit, pathogen infection, CO_2_, ozone, and darkness (Mustilli et al., 2002; Yoshida et al., 2002; Melotto et al., 2006; Merilo et al., 2013). However, there is significant functional redundancy between the three kinases. The Arabidopsis triple knockout mutant *snrk2.2/snrk2.3/snrk2.6* is extremely insensitive to ABA (much more than the single or double knockout mutants), exhibits severely impaired seed development and dormancy, and is oversensitive to water scarcity due to disruption of stomatal closure and down-regulation of ABA-and water stress-induced genes (Fujii and Zhu, 2009; Fujita et al., 2009; Nakashima et al., 2009). SnRK2s, which are either not or only weakly activated in response to ABA, are also involved in regulation of plant responses to abiotic stresses (Umezawa et al., 2004; Mizoguchi et al., 2010; Fujii et al., 2011; McLoughlin et al., 2012; Kulik et al., 2012; Soma et al., 2017).

The SnRK2 kinases are activated transiently in plant cells in response to environmental signals, and otherwise are maintained in inactive states. The best-known negative regulators of SnRK2s are protein phosphatases (Umezawa et al., 2009; Vlad et al., 2009; Hou et al., 2016; Krzywińska et al., 2016). Clade A phosphoprotein phosphatases 2C (PP2Cs) have been identified as major regulators of ABA-activated SnRK2s (Umezawa et al., 2009; Vlad et al., 2009; and reviews: Hubbard et al., 2010; Fujita et al., 2011; Nakashima and Yamaguchi-Shinozaki, 2013; Yoshida et al., 2015). Functional and structural studies showed that PP2Cs hold SnRK2s in an inactive state via a two-step inhibition mechanism (Soon et al., 2012; Zhou et al., 2012; Ng et al., 2014): specific Ser/Thr residues in the kinase activation loop are dephosphorylated, and a physical interaction between the kinase activation loop and the phosphatase active site persists to additionally block the kinase activity. Those results suggested that activity modulation is controlled not only by the phosphorylation state of SnRK2s but also by specific protein-protein interactions.

A few years ago, we identified and partially characterized another inhibitor of SnRK2 kinases and consequently of ABA signaling, SnRK2-interacting Calcium Sensor (SCS) (Bucholc et al., 2011). SCS provides the link between SnRK2s and calcium signaling pathways.

Calcium ions are ubiquitous second messengers that play pivotal roles in plant response to a number of external signals, inducing and regulating plant development, and responding to biotic and abiotic stresses. Several hundreds of plant proteins that potentially bind calcium have been identified; it was estimated that the Arabidopsis genome encodes about 250 EF-hand- or putative EF-hand-containing proteins (Day et al., 2002; Reddy and Reddy, 2004). Arabidopsis, presumably along with other plants, contain many more proteins with canonical and non-canonical EF-hand (“EF-hand-like”) motifs than other organisms. In plants, non-canonical EF-hand motifs are especially abundant (Day et al., 2002). Only a small fraction of them has been characterized. Numerous proteins with putative EF-hand-like motifs are involved in signal transduction; most probably they evolved to sense different calcium levels. Many of them contain four EF-hand sequences, with variable degrees of conservation of canonical EF-hand calcium binding motifs. This group comprises both sensor responder proteins (activated directly upon calcium binding and transmitting the signal further), as well as sensor relay proteins, which do not have enzymatic activity; upon Ca^2+^ binding, they undergo conformational changes and trigger activation or deactivation of their cellular partners. The best examples of sensor responder proteins are calcium dependent protein kinases (CDPKs), usually with four calmodulin-like EF-hand motifs and a Ser/Thr protein kinase domain that is activated upon calcium binding (review: Schulz et al., 2013). Sensor relay proteins involved in plant signaling constitute two main families: calmodulins (CaMs) and calcineurin B-like (CBL) proteins. Like CDPKs, each of them has four EF-hand or EF-hand-like motifs. Some CBLs harbor one or two canonical EF-hand motifs; most, however, have only EF-hand-like motifs (Batistič and Kudla, 2009; Sanchez-Barrena et al., 2013). The variety in calcium binding motif sequences determines the diversity in sensor proteins needed for sensing various calcium signatures, and finally to achieve the response specificity. However, for most CBLs and other predicted calcium binding proteins the actual calcium binding properties have not been characterized.

SCSs share several similar features with CBL proteins. According to the Prosite prediction of properties based on sequence, the *Arabidopsis thaliana* SCS protein (At4g38810) described by us earlier (Bucholc et al., 2011) contains four peptide sequences resembling EF-hand motifs, of which only one possesses all residues that define a canonical EF-hand. The other three are quite distinctly non-canonical and may be considered EF-hand-like. Also like CBLs, SCS is sensor relay protein involved in signal transduction in plants via interactions with SNF1-related protein kinases (SnRKs), unique to the plant kingdom. While SCSs interact with SnRK2s, CBLs interact with the SnRK3 kinases, also known as CBL-interacting protein kinases (CIPKs) or protein kinases related to SOS2 (PKSs). Both CBLs and SCSs regulate kinase activity, but in opposite ways, as CBLs activate CIPKs (for review see Batistič and Kudla, 2009; Luan, 2009; Batistič and Kudla, 2012) whereas SCSs inhibit SnRK2s.

Information provided by The Arabidopsis Information Resource (TAIR) indicates that in Arabidopsis two forms of AtSCS may exist, a longer one with 375 aa (denoted here as AtSCS-A, as described previously (Bucholc et al., 2011), and its shorter version with 265 aa (AtSCS-B) corresponding to the 111-375 aa fragment of AtSCS-A, in which the classical EF-hand motif is missing (Fig. 1A and 1B). *In silico* analysis of the *AtSCS* gene (At4g38810) sequence indicates that expression of both forms is possible due to the alternative transcription start sites (TSS); one promoter region is located upstream of ATG codon of *AtSCS-A*, and the second one responsible for *AtSCS-B* transcription within first *AtSCS-A* intron (Fig 1A). Alternative mRNA transcription starting, like alternative mRNA processing, is a well known regulatory process in eukaryotic organisms, including plants, which expands the genome’s coding capacity and generates protein variation (Tanaka et al., 2009; de Klerk and ’t Hoen, 2015). The result of alternative TSS are usually isoforms of proteins, which differ in their function, stability, localization and/or expression levels.

**Figure 1.**
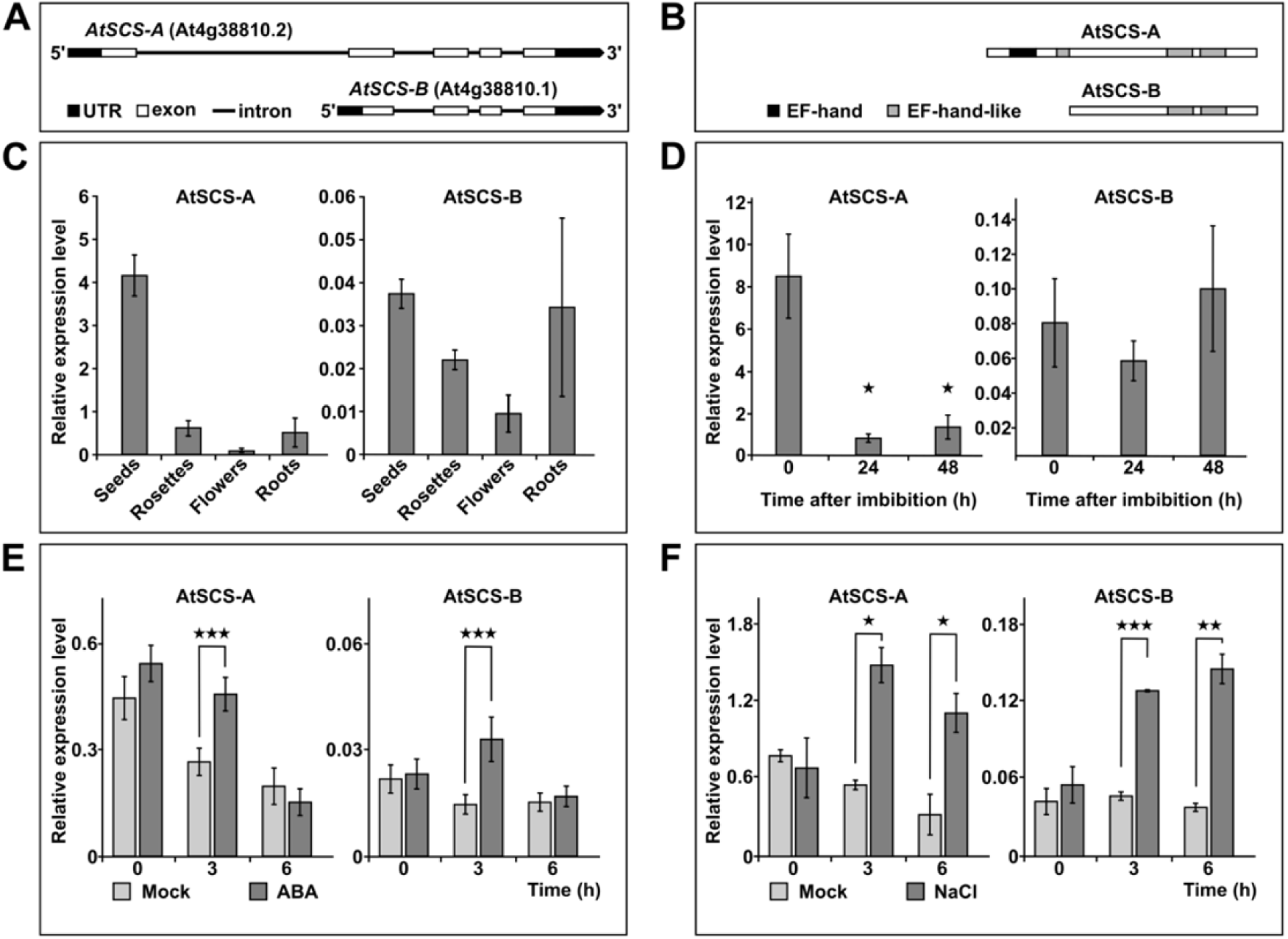
The expression of AtSCS-A and AtSCS-B varies across plant organs and in plant response to ABA or NaCl treatment. Alternative isoforms of AtSCS (At4g38810), AtSCS-A and AtSCS-B, predicted at transcript (A) and protein (B) levels. The prediction was done based on TAIR plant promoter database (PlantPromoterDB) and Prosite (prediction of EF-hand motifs). Quantitative RT-PCR analysis of *AtSCS-A* and *AtSCS-B* transcript levels in different organs (C) in seeds during germination (D), in 2-week-old seedlings exposed to 10 μM ABA (E) or 150 mM NaCl (F). Figures E and F - expression levels in plants exposed to ABA or NaCl (dark grey), whereas in control plants, mock treatment (light grey). Data represent means of triplicate biological repeats, and the error bars indicate SD. For statistical analysis a two-tailed *t-*test in Microsoft Office Excel was applied. The asterisks indicate significant difference from the wild type (*P < 0.05; **P < 0.01, ***P < 0.001).

In the present studies we confirmed the expression of both forms of AtSCS in Arabidopsis and showed significant differences in their expression profiles in plant organs and in response to abiotic stress. We compared both proteins with respect to Ca^2+^-binding properties, conformational dynamics with and without Ca^2+^, and regulation of the kinase activity *in vitro* and *in vivo*. Analysis of transgenic plants expressing *35S::AtSCS-A-c-myc* or *35S::AtSCS-B-c-myc* in the *scs-1* knockout background showed that both forms inhibit the ABA-responsive SnRK2s and, as a consequence, ABA signaling. The results show that both isoforms play a role in the regulation of the plant response to dehydration, analogous to that of the A-clade PP2Cs. Moreover, our results provide novel biochemical and biophysical data on EF-hand–like motifs in plant proteins.

## RESULTS

### AtSCS-A and AtSCS-B are Differently Expressed in Plant Tissues and in Arabidopsis Seedlings Subjected to ABA or Salt Stress

To verify that two forms of AtSCS are expressed in Arabidopsis, we analyzed mRNA level of *AtSCS-A* and *AtSCS-B* in various plant organs (seeds, rosettes, flowers, and roots). Moreover, since the functional analysis of AtSCS done previously by a reverse genetic approach indicated that AtSCS is involved in ABA signaling (Bucholc et al., 2011), we monitored *AtSCS-A* and *AtSCS-B* expression in seedlings exposed to ABA (10 μM) or salt stress (150 mM NaCl). To eliminate the potential influence of the circadian clock on the transcript levels, at each time point the transcript level of *AtSCS-A* and *AtSCS-B* was also monitored in plants not exposed to the stressor or ABA. The highest level of *AtSCS-A* transcript was observed in dry seeds, and the lowest in flowers (Fig. 1C and Supplemental Fig. S1). *AtSCS-B* expression was the highest in seeds and in roots, however, as it is shown in Fig. 1C and Fig. S1 *AtSCS-B* was expressed at a much lower level than *AtSCS-A* in all organs studied; in dry seeds expression of *AtSCS-A* was about 100 times higher than that of *AtSCS-B.* During seed imbibition the transcript level of *AtSCS-A* rapidly declined; after 24 h of imbibition the expression was about 10 times lower than that in dry seeds; in contrast, the transcript level of *AtSCS-B* did not change significantly during imbibition (Fig. 1D).

Analysis of *AtSCS-A* and *AtSCS-B* expression in seedlings exposed to 10 μM ABA showed transient increases (1.7- and 2.3-fold, respectively, at 3 hours) over levels in seedlings not exposed to ABA, with the differences disappearing by 6 hours after treatment (Fig. 1E). Salinity stress (150 mM NaCl) increased expression levels persistently (observed at both 3 and 6 h after treatment), to levels about 3 to 4-fold higher for both isoforms (Fig. 1F). It should be noted that only AtSCS-A expression undergoes changes during diurnal rhythm.

### AtSCS-B Interacts with Members of Group 2 and 3 of the SnRK2 Family

To determine whether AtSCS-B, like AtSCS-A, interacts with SnRK2s, we used a yeast two-hybrid approach. cDNA encoding AtSCS-B was fused in-frame to cDNA encoding the Gal4 activation domain in pGAD424 yeast expression vector, and cDNA encoding each of SnRK2s studied was fused to the Gal4 DNA-binding domain in pGBT9 vector. Using parallel constructions, we also studied interactions between AtSCS-A and SnRK2s. Those can be considered to be positive controls, since interactions between SnRK2s and AtSCS-A have been established previously (Bucholc et al., 2011). The results revealed that AtSCS-B interacts with SnRK2s from group 3 (SnRK2.2, SnRK2.3, and SnRK2.6) and from group 2 (SnRK2.7 and SnRK2.8), but does not interact with ABA-non-activated SnRK2 kinases from group 1 (SnRK2.1, SnRK2.4, SnRK2.5 and SnRK2.10) or with SnRK2.9 (Fig. 2A). The results suggest that AtSCS-B is a cellular regulator of members of groups 2 and 3 but not group 1 of the SnRK2 family. In contrast, AtSCS-A does not discriminate between these kinases in respect to the interaction. The only Arabidopsis SnRK2 that does not bind to either form of AtSCS is SnRK2.9.

**Figure 2.**
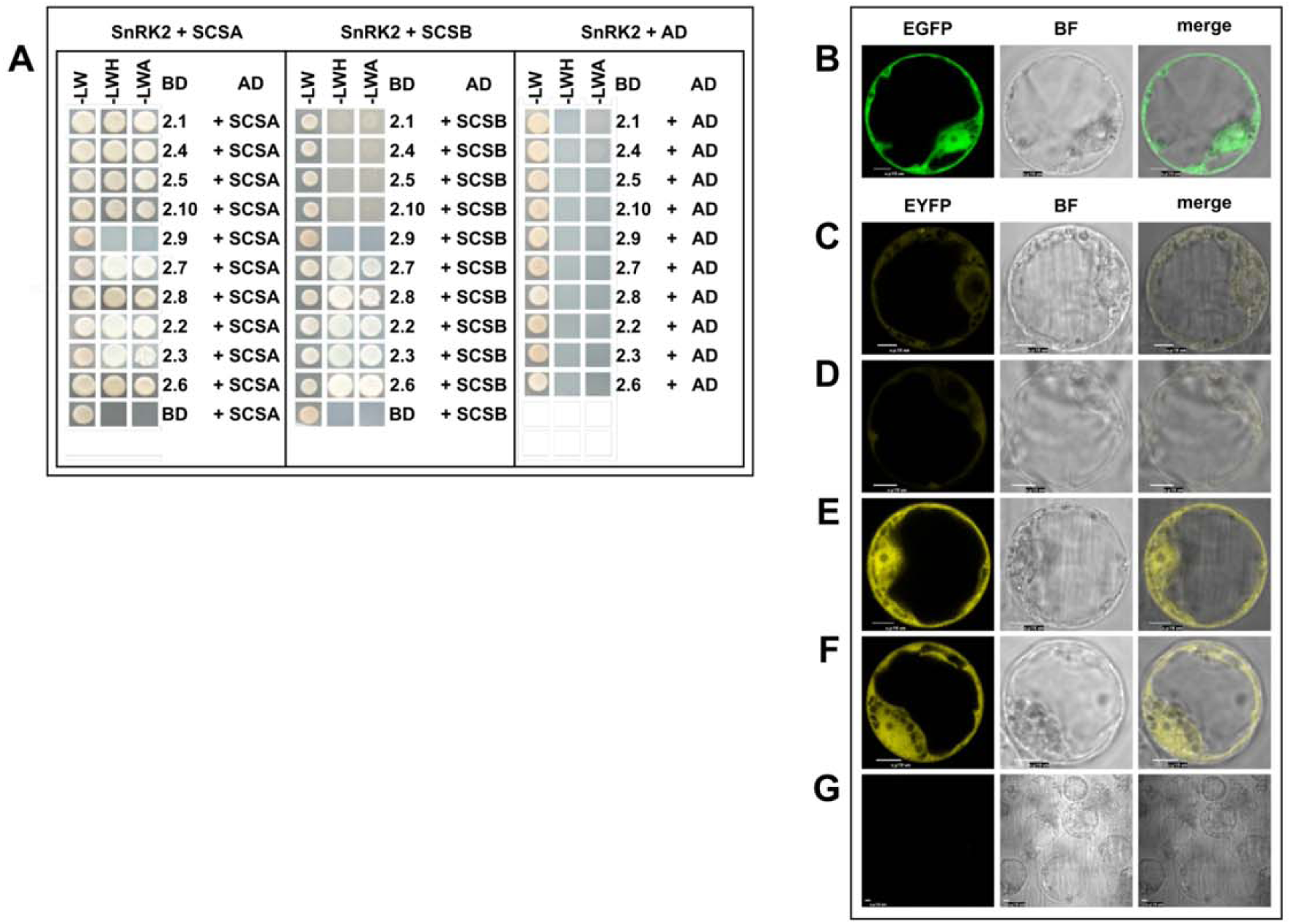
AtSCS-B interacts preferentially with kinases that belong to group 2 and 3 of the SnRK2 family. Interactions between Arabidopsis SnRK2s and AtSCS-B, or AtSCS-A as a control, were analyzed by a yeast two-hybrid assay (A), as described in Bucholc et al., 2011. Yeast transformed with a construct with cDNA encoding one of the analyzed SnRK2 and complementary empty vector (BD-SnRK2+AD), or a construct with AtSCS-B or AtSCS-A and the other empty vector (BD+AD-AtSCS) were used as controls. The growth of yeast expressing the indicated constructs was monitored on selective media: without Leu and Trp (-LW); without Leu, Trp and His and with 8mM AT (-LWH); without Leu, Trp and Ade (–LWA). AD, Gal4 activation domain; BD, Gal4 binding domain. The subcellular localization of AtSCS-B was analyzed in Arabidopsis protoplasts (B), as described in Bucholc et al., 2011. Protoplasts isolated from the T87 Arabidopsis cell line were transiently transformed with plasmid encoding AtSCS-B-EGFP and its localization was analyzed by confocal microscopy. Interaction between AtSCS-B and SnRK2s *in planta* was analyzed by BiFC assay. T87 protoplasts were transiently co-transformed with pairs of plasmids encoding: AtSCS-B-cEYFP and nEYFP-SnRK2.4 (C), AtSCS-B-cEYFP and nEYFP-SnRK2.10 (D), AtSCS-B-cEYFP and nEYFP-SnRK2.6 (E), AtSCS-B-cEYFP and nEYFP-SnRK2.8 (F). The binding led to reconstitution of functional YFP from chimeric proteins bearing non-florescent halves of YFP. For negative control, AtSCS-B-cEYFP was co-expressed with AtSCS-B-nEYFP (G). Scale bar = 10 µm; BF, bright field. The data shown here represent one of three independent experiments, all with similar results.

The interactions between AtSCS-B and the selected SnRK2s were verified by bimolecular fluorescence complementation (BiFC) assays. The proteins were expressed in Arabidopsis protoplasts as described in Bucholc et al. (2011). First, we analyzed the subcellular localization of AtSCS-B. The protein was produced as a fusion with EGFP using the pSAT6-EGFP-N1 or pSAT6-EGFP-C1 vector. As with AtSCS-A (Bucholc et al., 2011), AtSCS-B localized to the nucleus and cytoplasm (Fig. 2B). In a BiFC assay we analyzed interactions between AtSCS-B and SnRK2.4, SnRK2.10, SnRK2.6, and SnRK2.8 (Fig. 2C-2F). The kinases and AtSCS-B were each fused to complementary non-fluorescent fragments of YFP and transiently produced in Arabidopsis protoplasts. The BiFC assays showed that SnRK2.6 (from group 3) and SnRK2.8 (from group 2) interact with the AtSCS-B *in planta*. The interactions occur both in the cytoplasm and nucleus. SnRK2.4 and SnRK2.10 (from group 1) did not interact with AtSCS-B (in agreement with two-hybrid assay) or they interact very weakly, as the YFP signal is much weaker for these two kinases than that observed for SnRK2.6 or SnRK2.8. This supports the view that AtSCS-B is rather not a cellular regulator of ABA-non-activated SnRK2 kinases. A very low fluorescence signal detectable in the negative control samples was much weaker than YFP signal in BiFC samples (Fig. 2G). A comparison with AtSCS-A should be noted; as published previously, AtSCS-A interacts with all SnRK2 kinases studied, but exclusively in the cytoplasm (Bucholc et al., 2011).

### AtSCS-B Inhibits SnRK2 Activity in Calcium-Independent Manner

Our previous results showed that AtSCS-A inhibits SnRK2 activity only in the presence of calcium ions (Bucholc et al., 2011). To check whether AtSCS-B inhibition of SnRK2 activity is similarly calcium dependent we monitored the *in vitro* activity of SnRK2.6 and SnRK2.8 in the presence of increasing amounts of purified recombinant AtSCS-B, with and without calcium ions in the reaction mixture. The kinase activity was analyzed using MBP as a substrate. The results showed that AtSCS-B, in contrast to AtSCS-A, inhibits the SnRK2 activity both in the presence and in the absence of calcium ions (in the presence of EGTA) (Fig. 3A and 3B).

**Figure 3.**
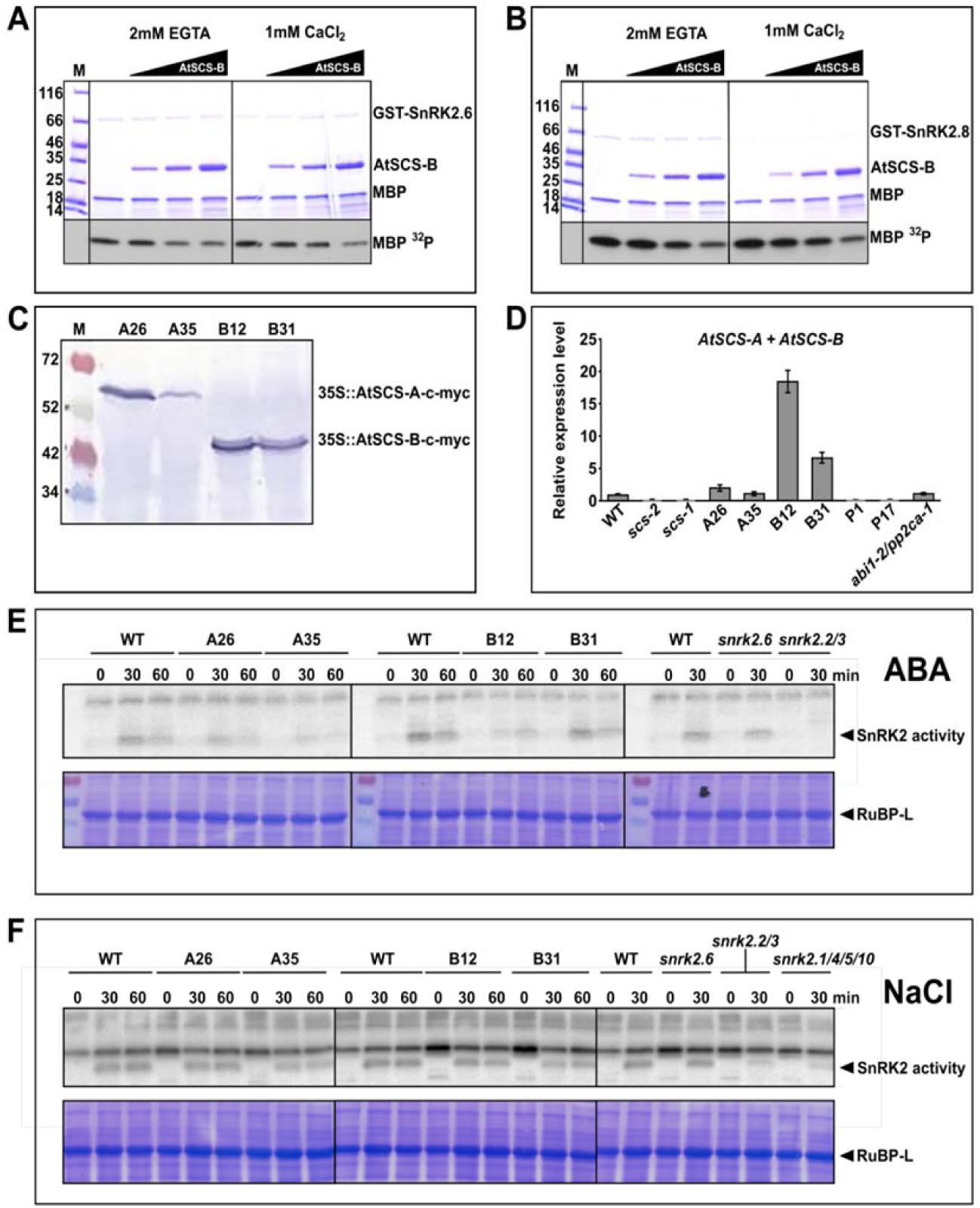
AtSCS-A and AtSCS-B inhibit the SnRK2 activity in vitro and in vivo. AtSCS-B inhibition of SnRK2.6 (A) and SnRK2.8 (B) is calcium independent. SnRK2.6, SnRK2.8, and AtSCS-B were expressed in *E.coli* and the kinase activity was measured in the presence of increasing amounts of AtSCS-B (0, 40, 80 and 160 ng per µL) without or with Ca^2+^ (2 mM EGTA or 1 mM CaCl_2_, respectively) in the reaction mixture. The kinase activity was monitored using MBP and [γ^32^P]ATP as substrates. Reaction products were separated by SDS-PAGE and MBP phosphorylation was determined by autoradiography. The data represent one of three independent experiments showing similar results. The expression of AtSCS-A and AtSCS-B in seedlings of homozygous transgenic lines 35S::AtSCS-A-c-myc (A26 and A35) and 35S::AtSCS-B-c-myc (B12 and B31) was measured at protein (C) and transcript (D) levels. The production of AtSCS-A-c-myc and AtSCS-B-c-myc proteins was monitored in seedlings of the transgenic plants by Western blotting using anti-c-myc antibodies. *AtSCS* mRNA level was monitored in the transgenic plants expressing *AtSCS-A-c-myc* or *AtSCS-B-c-myc*, or *c-myc* (vector control: P1, P17), the *scs-1* and *scs-2* mutants, as well in the WT plants by quantitative RT-PCR analysis. Data represent means of triplicate biological repeats, and the error bars indicate SD. Analysis of the SnRK2 activity in 2-week-old seedlings of the WT, transgenic plants expressing *AtSCS-A-c-myc* or *AtSCS-B-c-myc*, and selected *snrk2* knockout mutants subjected to 100 μM ABA (E) or 350 mM NaCl (F) shows significant *in planta* inhibition by AtSCS isoforms. The kinase activity was monitored by in-gel-kinase activity assay using as substrate GST-ABF2 (G73-Q120) or MBP, respectively. The representative results from one of three independent experiments are shown.

### Both AtSCS-A and AtSCS-B Inhibit the SnRK2 Activity *in planta*

To investigate the involvement of AtSCS-A and AtSCS-B in the regulation of SnRK2 activity *in vivo* we obtained homozygous transgenic Col-0 Arabidopsis plants expressing *AtSCS-A-c-myc* (5 independent lines) or *AtSCS-B-c-myc* (3 independent lines) under control of the 35S promoter, as well as plants expressing an empty vector, in an *scs-1* knockout background deficient in both AtSCS forms. For the studies we selected two transgenic lines 35S::AtSCS-A-c-myc (A26 and A35), 35S::AtSCS-B-c-myc (B12 and B31), with the highest expression of the transgene (Fig. 3C, 3D, and Supplemental Fig. S2). We analyzed the kinase activity phosphorylating ABF2 (Gly73-Gln120) peptide (in the case of plants exposed to ABA) or myelin basic protein (MBP, in the case of plants exposed to NaCl) in 2-week-old seedlings of A26, A35, B12, and B31 lines, and the wild type Arabidopsis (WT). The activity was analyzed before and after treatment with 100 μM ABA (or 350 mM NaCl) by in-gel kinase activity assay according to Wang and Zhu (2016). The SnRK2 activity induced in response to ABA was significantly lower in both 35S::AtSCS-A-c-myc lines (A26 and A35) and in one of 35S::AtSCS-B-c-myc lines (B12) compared to WT plants (Fig. 3E). In the B31 line, with its lower expression of AtSCS-B-c-myc compared to B12 line, the ABA-induced SnRK2 activity was similar to that observed in the WT plants. These results indicate that both AtSCS-A and AtSCS-B inhibit the kinase activity induced by ABA treatment; however, the inhibition is stronger in the presence of AtSCS-A. The analysis of the kinase activity in the same Arabidopsis lines treated with NaCl showed only very weak, if any, inhibition of the SnRK2 activity by AtSCSs (Fig 3F). In summary, these results indicate that both forms of AtSCS are able to inhibit the SnRK2s *in vivo*, especially those kinases which are involved in ABA signaling.

### AtSCS-A and AtSCS-B Have an Impact on the Plant Sensitivity to Dehydration

Our results showed that expression of *AtSCS-A* and *AtSCS–B* is induced in the response to ABA (Fig.1E). Moreover, the activity of ABA-responsive kinases in Arabidopsis seedlings overexpressing AtSCS-A or AtSCS-B exposed to exogenous ABA is significantly reduced in comparison to that observed in the WT plants (Fig. 3E). Since the ABA-responsive SnRK2s are key regulators of the plant response to dehydration, we studied the impact of AtSCSs on the plant response to water deficit. We analyzed the survival rate of the Col-0 WT, the *scs* knockout mutants (lines *scs-2* and vector control P1, 35S:c-myc in *scs-1* background), 35S::AtSCS-A-c-myc (A26 and A35 lines) and 35S::AtSCS-B-c-myc (B12 and B31lines) transgenic plants under drought conditions (the watering was withdrawn for 14 days) and after re-watering. In our assay we also included an *abi1-2/pp2ca-1* mutant deficient in two clade A PP2C phosphates, ABI1 and PP2CA, well known inhibitors of ABA-dependent SnRK2s and ABA signaling (Umezawa et al., 2009; Vlad et al., 2009; Rubio et al., 2009), as a control. Our results confirmed previously published data (Rubio et al., 2009) that the *abi1-2/pp2ca-1* mutant exhibits enhanced resistance to drought stress and showed that the *scs-2* mutant and P1 line similarly as *abi1-2/pp2ca-1* were more resistant to dehydration than all other lines studied (Fig. 4A and 4B). The data indicated similar regulation of the response to dehydration of AtSCSs and the clade A PP2Cs. The results also showed that expression of *AtSCS-A* (lines A26 and A35) or *AtSCS-B* (especially line B12 with a higher level of AtSCS-B) alone only partially complement the phenotype of the *scs* mutant, which suggests that both forms of SCS are needed for the full complementation.

**Figure 4.**
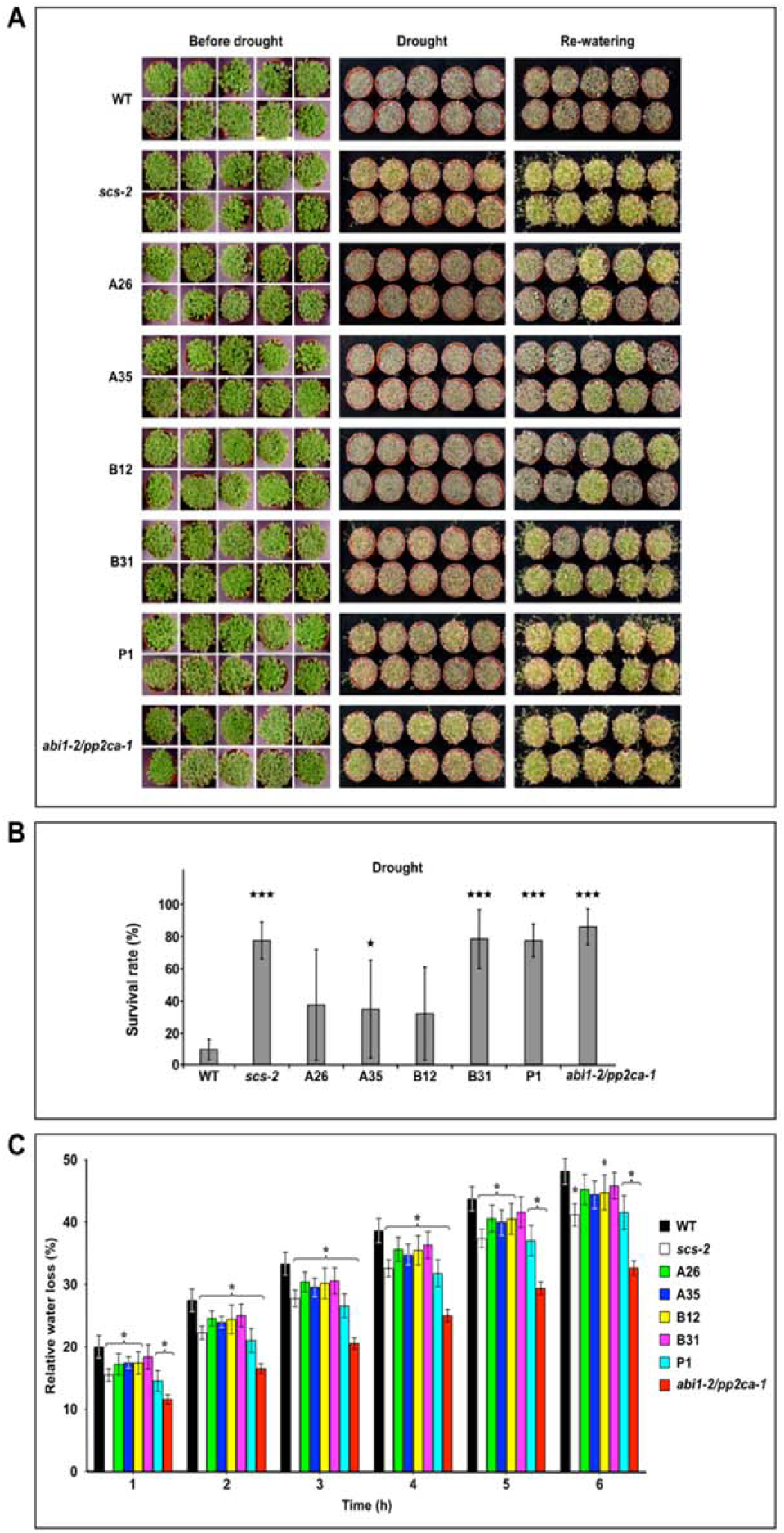
AtSCS-A and AtSCS-B Regulate the Response of Arabidopsis to Water Deficit. The drought survival rate test (A). The Arabidopsis plants were grown in pots for 17 days under long day conditions and for an additional 2 weeks without watering. The pictures were taken before watering was stopped (before drought), after two weeks without water (drought), and two days after re-watering (re-watering). Ten pots with approximately 50 plants per pot for each line per experiment were used. Representative plants are presented. The drought survival rate (B) was estimated by analysis of at least 1000 plants per each line. For the statistical analysis a t-test was applied. The asterisks indicate significant differences from the wild type (*P < 0.05; **P < 0.01, ***P < 0.001). The average values ± SE are shown. Water loss from detached Arabidopsis rosettes (C). Rates of water loss from the whole rosettes of six-week-old plants of wild type and different mutant lines were measured at the time points indicated. Finally, the rosettes were dried at 70°C overnight and weighed. The cut rosette water loss (CRWL) was calculated. The representative results from one of four independent experiments are shown. Eight plants were used for each line per experiment. For the statistical analysis, a t-test was applied. The asterisks indicate significant differences from the wild type (*P < 0.05; **P < 0.01, ***P < 0.001). The average values ± SE are shown.

Additionally, we measured water loss in detached rosettes of 6-week-old plants of all the lines listed above. The water loss was lower in rosettes of the *scs* mutants (*scs2* and P1, vector control in *scs1* background) than in other lines (Fig. 4C), which is in agreement with the result of drought survival test (Fig. 4A and 4B).

Using Arabidopsis lines with differing the AtSCS-A or AtSCS-B levels we also investigated the involvement of AtSCSs in regulation of the expression of ABA-induced stress-related genes. For the analysis we chose two genes regulated by SnRK2s in an ABA-dependent manner, *Rab18* and *RD29B*. We did not observe any significant differences in the expression of genes studied in response to ABA between lines studied (Supplemental Fig S3) indicating that SCSs are rather not involved in the regulation of gene expression. This result is consistent with our data showing that SCSs interact with SnRK2s mainly in the cytoplasm; AtSCS-A interacts exclusively in the cytoplasm (Bucholc et al., 2011, Fig.2B), whereas AtSCS-B in the cytoplasm and in the nucleus. However, we cannot exclude the possibility that transiently overexpressed AtSCS-B-GFP/YFP is passively diffused to the nucleus (that is why we observe its presence both in the cytoplasm and the nucleus).

### AtSCS-B Binds Ca^2+^ Without a Substantial Effect on the Protein Conformation

In order to compare calcium-binding properties of AtSCS-B to that of AtSCS-A described earlier (Bucholc et al., 2011), we monitored changes in fluorescence of AtSCS-B and AtSCS-A (as a control) accompanying binding of calcium. The proteins were excited at 280 nm (monitoring the fluorescence of all fluorophores) or at 295 nm (monitoring the fluorescence of a Trp residue). AtSCS-B protein has three fluorophores: Trp72, Tyr29 and Tyr168 (corresponding to Trp182, Tyr139 and Tyr278 in AtSCS-A). The protein fluorescence spectrum is dominated by the broad tryptophan fluorescence band, and the tyrosine fluorescence band is apparent as well. The binding of calcium caused small changes: the small intensity decrease and very small ‘red shift’ of the maximum, λmax (apo) = 342.5 nm → λmax (*holo*) = 343 nm (for the fluorescence excitation at 280 nm) and λmax (*apo*) = 350 nm → λmax (*holo*) = 351nm (excitation at 295 nm) (Fig. 5A). Also, the small increase of the intensity was observed for the fluorescence band of the tyrosine residue(s), near 305 nm. The fluorescence spectrum of AtSCS-B suggests that the tryptophan residue is buried and inaccessible to an energy transfer from the tyrosine residue(s), and located in an environment that quenches its fluorescence strongly. The calcium binding entails a subtle rearrangement of Trp72 environment, linked with the small red shift of Trp72 fluorescence (at 295 nm excitation) (Fig. 5A).

**Figure 5.**
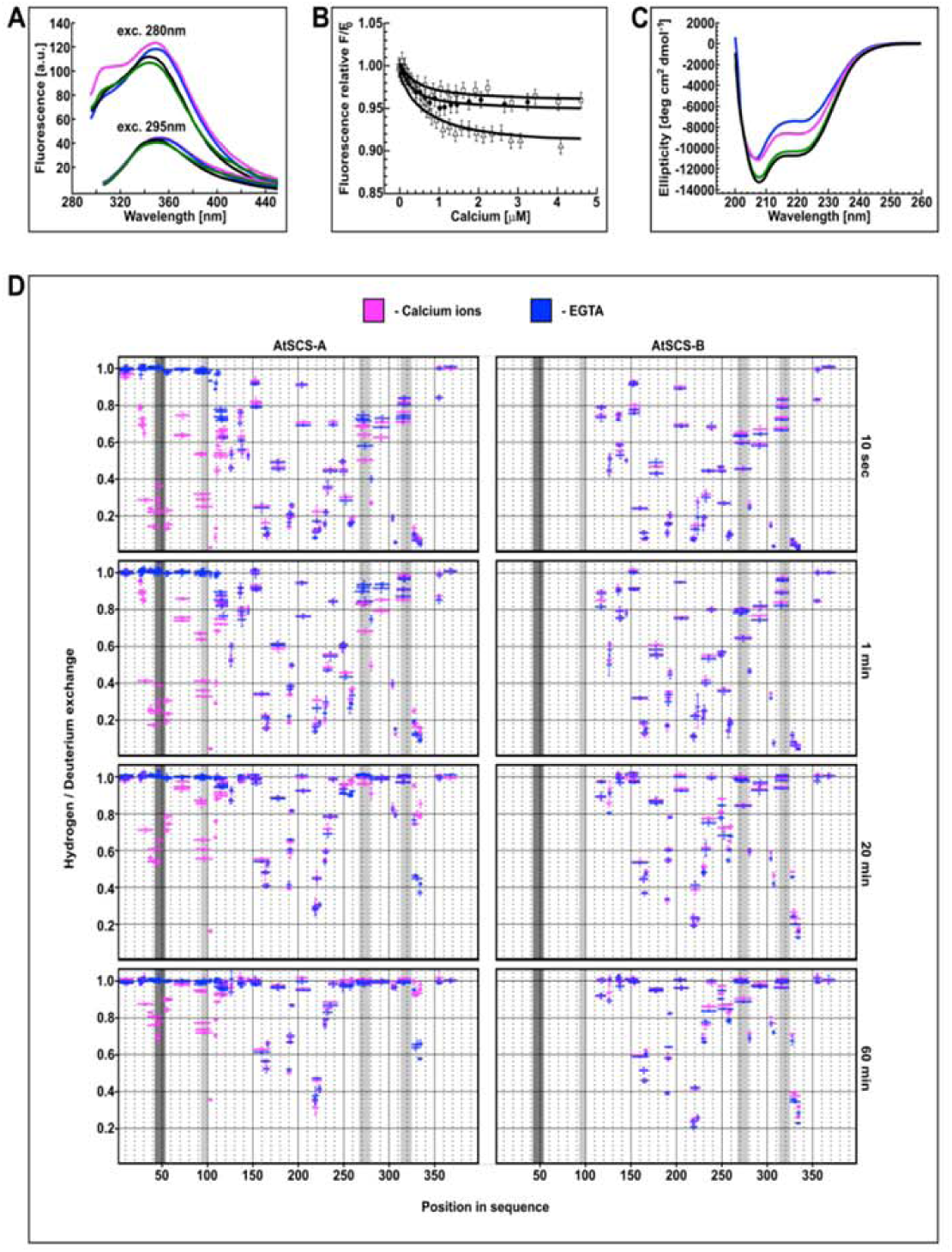
AtSCS-B Binds Ca2+ Without a Substantial Effect on the Protein Conformation. The fluorescence spectra of AtSCS-B and AtSCS-A proteins (A). The fluorescence spectra of 1.2 µM solution of AtSCS-A or AtSCS-B protein, excited at 280nm (upper curves) or at 295nm (bottom curves). apo AtSCS-A protein (blue line), apo AtSCS-B protein (black line), holo AtSCS-A (magenta line), and holo AtSCS-B (green line) are presented. The fluorescence spectra represent one of three independent measurements. The fluorescence titration curves of AtSCS-B protein by calcium ions (B). The fluorescence excitation and the emission were at 280 and 342 nm (open triangles) or 280 and 340 nm (black circles, open squares), respectively. The calcium ions titration of 1.5 µM solutions of AtSCS-B as well as the experimental errors and the theoretical curves, calculated for a single-site ligand binding model (solid lines), are shown. Conformational changes of AtSCS-A and AtSCS-B proteins upon Ca^2+^binding. (C) CD spectra of 2 μM solutions of AtSCS-B or AtSCS-A with 1 mM Ca^2+^ (green and magenta lines, respectively) or without Ca^2+^ (100 μM EDTA; blue for apo AtSCS-A and black for apo-AtSCS-B). Changes of conformational dynamics of AtSCS-A and AtSCS-B upon Ca^2+^ binding. (D) H/D exchange in AtSCS-A (left panels) and AtSCS-B (right panels) in the presence (magenta) or in the absence of Ca^2+^ (EGTA, blue). The proteins were incubated in D_2_O buffer for various times (10 sec, 1 min, 20 min, 60 min) and the extent of H/D exchange was determined by mass spectrometry. The horizontal red and blue bars indicate individual peptides identified by mass spectrometry. The X-axis indicates their position in the amino acid sequence of the AtSCS-A. The Y-axis shows relative deuterium uptake calculated as described in Material and Methods; the value 1 represents maximal deuteration level, meaning that all hydrogens in amide bonds of particular peptide were exchanged to deuterium. Error bars are standard deviations from at least three independent experiments. Dark grey – position of calcium binding loops of the canonical EF-hand motif, light grey – positions calcium binding loops of potential non-canonical EF-hand motifs.

In parallel, analogous experiments were performed for AtSCS-A protein, as was also studied previously (Bucholc et al., 2011). AtSCS-A protein is longer than AtSCS-B by 110 amino acid residues and richer by one tyrosine residue, Tyr80. The protein is also more sensitive to calcium ions due to the presence of the canonical EF-hand motif. The fluorescence spectra of the AtSCS-A, for *apo* form of the protein, are similar to those of AtSCS-B (Fig. 5A). The most significant fluorescence changes after calcium binding were observed as an increase in the intensity of the tyrosine fluorescence band (Fig. 5A). The contribution of tryptophan residue to those changes is small. The fluorescence spectra excited at 295 nm, are almost the same for the calcium-free and the calcium-bound protein (Fig. 5A, bottom curves), supporting the conclusion that the conformation of the protein in the environment residues Trp182 is only slightly altered by calcium ions. In contrast, the conformation of the protein in the environment of the one or more of the tyrosine residues changes dramatically. Tyr80 present in the N-terminal fragment of AtSCS-A (absent in AtSCS-B protein) seems responsible for these changes. Its fluorescence is influenced most probably by the rearrangement of the canonical EF-hand motif upon calcium binding.

The affinity for calcium of AtSCS-B protein was determined from the fluorescence titration experiments (Fig. 5B). The fluorescence, excited at 280 nm, was measured at 340 or 342 nm, in the fluorescence maximum. The maximal fluorescence intensity changes were small; they did not exceed 10% of the initial fluorescence of the protein (Fig. 5B). To the resulting titration curve, the function describing the binding of calcium ion [eq. (c), from “Experimental Procedures”] was fitted. The function is analogous with eq. 8 in Eftink (1997) describing single-site ligand binding. The calcium association constant K1 was estimated as 2.4(±1.0)×10^3^ M^−1^, as the average of three independent measurements. The value of K1=2.4×10^3^ M^−1^, corresponds to a dissociation constant of 0.4 mM.

The stoichiometry of the binding of calcium ions to AtSCS-B was determined from the fluorescence titration curves (Supplemental Fig. S4). The index n equal 1 was estimated based on the Hill equation describing the association of the ligand to the protein, used for the analysis of fluorescence measurements. The resulting formula, derived from the classic Hill equation, is as follows:

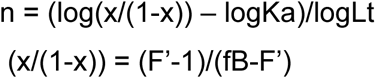

where: × – mole fraction of the complex protein-ligand, F’ - relative observed fluorescence (relative to the initial fluorescence without the ligand), fB - relative fluorescence of the complex protein-ligand, Lt - the total (free and bound) concentration of ligand, and Ka is the overall association constant of protein-ligand. This result indicates that AtSCS-B molecule most probably binds one calcium ion.

Structural changes of AtSCS-B upon calcium binding were analyzed by far-UV circular dichroism spectroscopy (CD). The binding of calcium ions by the AtSCS-B protein was accompanied by minor structural changes, seen as a decrease of negative CD signals (Fig. 5C) indicating very mild disruption of the protein structure in response to calcium binding, limited to some reduction of helix content or some helix reorientation. In parallel, as a control, the CD spectrum of AtSCS-A protein in the same conditions was analyzed (Fig. 5C). The values of residual molar ellipticity agreed well with the earlier data (Bucholc et al., 2011). The Ca^2+^-free AtSCS-A protein is 28% helical, and the contribution of helical structures in the Ca^2+^-bound protein is higher, but less than in AtSCS-B protein (both calcium-free and calcium-bound).

### Differences in Conformational Dynamics Between AtSCS-A and AtSCS-B upon Calcium Binding

The analysis of AtSCS proteins using fluorescence and CD spectroscopy methods revealed that the structure of AtSCS-A, but not AtSCS-B, undergoes significant change upon Ca^2+^ addition; they do not, however, indicate where the conformational changes occur. Therefore, in order to investigate the structure and conformational dynamics of AtSCS proteins in the absence and in presence of Ca^2+^ ions, hydrogen deuterium exchange monitored by mass spectrometry (HDX-MS) was applied. HDX-MS is an analytical technique that maps protein conformational dynamics in solution to specific regions of the protein. The approach applied in this study gave structural information with peptide resolution since the measured degree of exchange is an average over the lengths of the peptides resulting from pepsin digestion. This is especially informative, as to our knowledge no structure of AtSCS-A or AtSCS-B (or any similar proteins) has been determined by X-ray crystallography, Cryo-EM or NMR; none has been deposited in the PDB database.

The hydrogen deuterium exchange patterns along the AtSCS-A and AtSCS-B proteins were obtained for two conditions, one in the presence and in the absence of Ca^2+^ (in buffer with EGTA). Regions could be classified variously as stable or labile, and the stability of some of these regions was calcium dependent, as we describe next.

#### Conformational dynamics of AtSCS-A

The HDX-MS data show that AtSCS-A protein contains several regions of retarded exchange, independent of the presence of Ca^2+^ (Fig. 5D and Supplemental Fig. S5-S7). Peptides spanning 151-169, 187-196, 215-226 residues can be classified as the most protected (i.e., most stable) regions. These are thus regions involved in the hydrogen-bonding network that constitute the structural core of the molecule. There are also regions along the AtSCS-A sequence for which the level of HD exchange is very high, also independent of the presence of Ca^2+^ ions. Peptides spanning positions 1-19, 110-123, 133-140, 147-160, 197-214, 233-244, 307-325, 351-375 can be classified as the most labile regions. Highly dynamic regions in the inner parts of the SCS protein sequence are probably located within loops and extended turns which connect adjacent elements of the protein.

Some segments of AtSCS-A showed drastically different conformational dynamics depending on the presence of calcium ions. In the presence of EGTA (and absence of calcium), an N-terminal segment of AtSCS-A (region approximately from 20 to 110 amino acid) undergoes very fast HD exchange. This region exhibits no protection even at the lowest measured experimental time of 10 seconds labeling (Fig. 5D, Supplemental Fig. S5-S7), indicating a lack of secondary structure of this domain. The presence of Ca^2+^ ions dramatically changes the pattern of HD exchange in this region, strongly decreasing the uptake of deuterium in two segments. One of them spans the calcium-binding loop of classical EF-hand motif (fragment 42-53). Unexpectedly, significant changes in the deuterium uptake after Ca^2+^ addition occur also in the region of residues 85-112 (Fig. 5D, Supplemental Fig. S5-S7). In that part of the AtSCS-A, a Prosite motif search algorithm predicted the occurrence of the putative non-canonical EF-hand motif.

Some changes in levels of HD exchange upon Ca^2+^ binding also appear in the AtSCS-A segments 254--301 and 325-337 (Fig. 5D, Supplemental Fig. S6-S7). Two hypothetical EF-hand-like loops (positions: 267-278 and 313-324) are predicted within and close to these fragments. Here, several regions characterized by relatively stable structure were detected. Notably, the retarded exchange is observed in segments directly flanking the putative calcium binding motifs, indicating the presence of stable secondary structure elements there (positions: 245-263 and 325-337). The flanking regions seem to be structured both in the presence and in the absence of calcium ions, whereas the putative EF-hand-like calcium-binding loops exhibit quite high HD exchange levels independent of calcium ions (Fig. 5D and Supplemental Fig. S5 and S6) and can be classified as dynamic regions. Interestingly, the flexibility of fragment 262-301 is decreased when calcium is present (Fig. 5D, Supplemental Fig. S5 and S6). The opposite effect, i.e., destabilization in the presence of calcium, was observed for the region 325-337.

#### The structure of the common region is more stable in AtSCS-B than in AtSCS-A, both in the presence and in the absence of calcium ions

Patterns of HD exchange obtained in the same experimental conditions for AtSCS-A and AtSCS-B indicate that in general both forms do not differ much from each other in the region 110-250, which is common for both forms (Fig. 5D and Supplemental Fig. S5 and S7). There are however some differences, mainly near putative calcium binding loops (region 157-168 in AtSCS-B, corresponding to 267-278 in AtSCS-A, and 203-214 in AtSCS-B, corresponding to 313-324 in AtSCS-A. Strikingly, in the longer variant (AtSCS-A), regions flanking the predicted calcium-binding loops reveal lower stability than in the shorter variant, which lacks residues 1-110 (AtSCS-B). Thus, the native stability of the C-terminal fragment is disturbed in the structural context of the longer variant. This strongly indicates that the N-terminal domain, absent in AtSCS-B, is structurally coupled to the C-terminal part in AtSCS-A and destabilizes its structure. On the other hand, the HDX data show that in the presence of calcium the third predicted EF-hand-like calcium-binding loop (region 267-278) in AtSCS-A is stabilized in the presence of calcium and becomes similar to that of AtSCS-B. (Fig.5D and Supplemental Fig. S5 and S6)

### Models of AtSCS-A / AtSCS-B Structure

Due to the lack of relevant templates in the PDB, the 3D structure of AtSCS-A was modeled separately for two regions, defined by residue ranges of 1-175 and 211-330, respectively. For these protein fragments homology modeling was carried out based on the rationally selected subsets of PDB structures; as described in Methods section.

#### N-terminal domain of AtSCS-A

The best models for the fragment covering residues 1-175 were built using either 4MBE or 1SL8 PDB records as templates. However, in the hybrid model, obtained by the combination of fragments of 4MBE, 4OR9, 1SL8 and 4N5X, modeled structures were scored even better. Since the N-terminal fragment 1-26 was strongly divergent in all models, and was also found flexible in HDX experiments, the subsequent round of modeling was restricted to residues 26-175. The final hybrid model, based on the combination of structural motifs adopted from 2ZND, 1Y1X, 5B8I, 2TN4, 4OR9, scored higher than any of the single structure-based models. The fold of the modeled N-terminal domain shows two pairs of EF-hand-like motifs, the relative orientation of which visibly varies between individual models. However, in the majority of them, the ^42^**D**Q**D**E**D**G**K**LSVT**E**^53^ loop (Fig. 6A, magenta tube) adopts the canonical conformation characteristic for the calcium-loaded form of EF-hand, whereby calcium is coordinated by the side-chain carboxyl groups of Asp42, Asp44, Asp46 and Glu53, along with the carbonyl oxygen of Lys48 (magenta in Fig. 6A). It should be however mentioned that, according to the review on conservation of individual residues located at the Ca^2+^ binding loop (Halling et al., 2016) Gln, Glu, Lys, Leu, Val and Thr are rarely identified at positions 2, 4, 7, 8 and 11 of calmodulin EF-hand motifs (1.0, 1.1, 5.2, 1.8, 2.0 and 3.0 %, respectively). The sequences of the paired loop (^88^**T**H**G**S**Q**E**K**VSKT**E**^99^, blue in Fig. 6A) preclude calcium binding. Interestingly, the side chain of Lys94 located in this loop may compete with Ca^2+^ for the binding site, thus stabilizing an EF-hand fold for the *apo* form. Both findings suggest that the N-terminal domain of AtSCS-A binds a single Ca^2+^ cation with an affinity substantially lower than that of calmodulin, which confirms our previous data describing Ca^2+^-binding properties of AtSCS-A (Bucholc et al., 2011).

**Figure 6.**
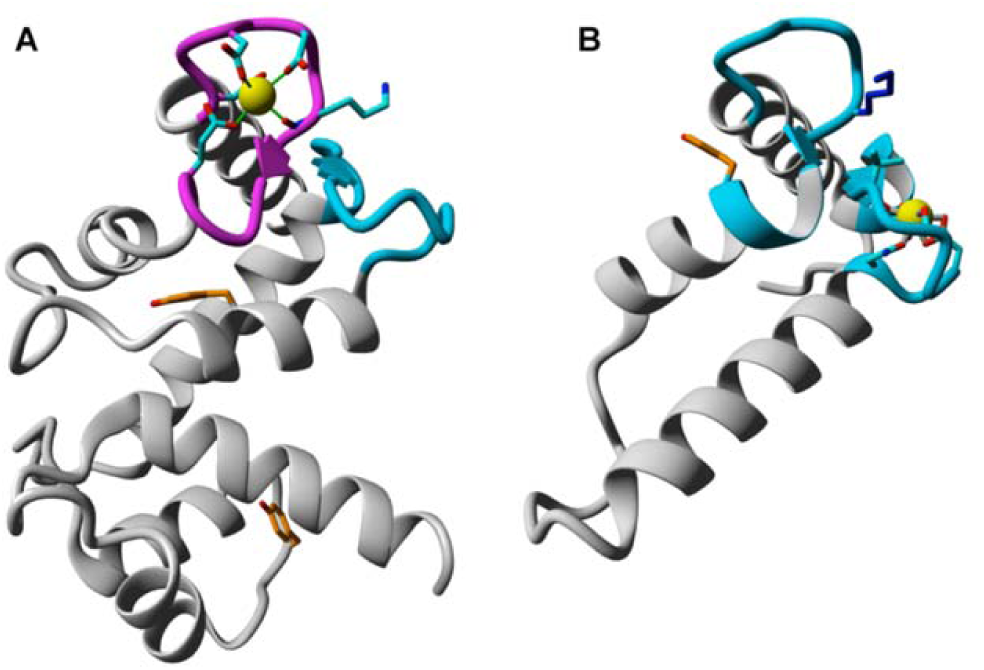
Modeled by homology structures of N-terminal (residues 26-175) (A) and C-terminal (residues 251-330) (B) parts of AtSCS-A. The ribbon representation follows the protein backbone. In magenta and blue are marked peptides predicted as canonical or non-canonical, respectively, EF-hand calcium binding loops. Residues putatively involved in calcium binding are denoted by sticks, yellow spheres represent calcium ions. Tyrosine residues are labeled by orange color.

Finally, it should be mentioned that both tyrosine residues (Tyr80 and Tyr139, orange in Fig. 6A) are buried, and fluorescence of Tyr80 should be affected by calcium binding, what is in agreement with our results of the analysis of AtSCS-A in respect of changes in fluorescence upon binding of calcium.

#### C-terminal domain of AtSCS-A / AtSCS-B

There were no appropriate templates for the C-terminal fragment (residues 331-378). The best model for the fragment defined by residues 211-330 was based on 1PRW, and no further improvement was achieved for the hybrid models. However, detailed inspection of the alternative highly scored models (e.g., based on 1QTX or 4AQR) clearly shown that only the fragment covering residues 251N-A330 could be reasonably modeled, while the homology-modeled structure of 211-250 critically depends on the template protein used. The subsequently built hybrid model for 251N-A330 was based on 3TZ1, with some parts improved using alternative models based on 1QTX, 1RFJ, 3WFN and 1PRW. The resulting structure represents pair of EF-hand-like motifs, one of which (^313^NG**D**D**G**NVVKEEE^324^) may display propensities toward binding of calcium via side-chain oxygen atoms of N313, D315 and E324, and backbone carbonyl groups of G317 and V319, the affinity of which must be expected much lower than that for N-terminal part of the protein. This loop is putatively paired with (^267^PK**D**RQGKVSKGY^278^), sequence of which preclude any interaction with calcium (Fig. 6B). It should be noted that side chain of K273 might interact with the ^313^NGDDGNVVKEEE^324^ loop, thus mimicking calcium-loaded EF-hand state.

Our HDX data indicate interplay between the N-terminal part of AtSCS-A (containing the canonical EF-hand motif) and the C-terminal part (containing only EF-hand-like motifs). Both domains seem to be structurally coupled in AtSCS-A, and strong structuring of the N-terminal region of the AtSCS-A upon Ca^2+^ binding affects this interaction. Calcium-induced structuring of the AtSCS-A N-terminal domain may influence the dynamics of distant parts, which are already well structured in the *apo* form (Fig. 5D). It is possible that the small changes observed in the C-terminal fragment of AtSCS-A upon binding of calcium do not indicate direct calcium binding to this domain, but may originate from an allosteric transmission caused by the calcium binding to the canonical site in the N-terminal domain. Fragment 325-337 FKKTMAEILGSIM, whose stability in AtSCS-B is nearly independent of the presence of Ca^2+^, is definitely less stable in AtSCS-A *apo*, and is even much more destabilized in AtSCS-A in the presence of calcium ions (Fig. 5D, Supplemental Fig. S6 and S7). In contrast, the region 278-283 YLRAVL (Supplemental Fig. S7 and S8) in AtSCS-A in the presence of Ca^2+^ becomes stabilized and at least in 1 min of exchange resembles this fragment within AtSCS-B.

According to the model, the first calcium-binding domain (canonical EF-hand motif) forms a classical helix-turn-helix structure with helices 26-40 and 57-63 (Fig. 6A). This model agrees well with the HDX data. In the presence of Ca^2+^, regions characterized by the strongest retardation of HD exchange cover the 28-62 fragment (representing canonical EF-hand motif), and additionally the 85-115 one, where the EF-hand-like motif is predicted. The helices of canonical EF-hand and non-canonical EF-hand exhibit similar HD exchange patterns, with a strongly retarded exchange in the presence of Ca^2+^ with the exception of the helix at 77-87, where HD exchange is much faster. The model predicts proximity of the canonical EF-hand motif and the non-canonical one. For canonical EF-hand motif the lowest level of HD exchange is observed within loop 41-56 constituting Ca^2+^ binding sites, for non-canonical EF-hand the lowest level of HD exchange refers to fragment 101-106 within helix 99-117 (Fig. 5D and Supplemental Fig. S7 and S8).

Modeling of the region 250-330 predicts the existence of four helices (252-256, 275-287, 297-313, 321-326) (Fig. 6B). Indeed, in the HDX experiment there is significant retardation in HD exchange within helix 252-256, 275-287, 297-313 and 321-326 (Fig. 5D and Supplemental Fig. S8). The structural model predicts direct contact between helices. HDX technique confirms the stability of regions above, consistent with the modeled interaction between adjacent helices, which should significantly stabilize the network of hydrogen bonds and create a connected and coherent structure (Fig. 6B).

## DISCUSSION

Depending on a variety of environmental cues, plants trigger appropriate responses from among diverse signaling pathways, activating specific defense mechanisms and adjusting plant development to new conditions. SnRK2 kinases play a key role in the regulation of the ABA-dependent development and responses to water deficit, as well as several other stresses. They are known as calcium-independent enzymes; however, several data indicate their interplay with calcium signaling pathways. Calcium-dependent kinases, CIPKs, together with SnRK2.6 phosphorylate and regulate activity of NADPH oxidase (respiratory burst oxidase homolog F, RbohF; Sirichandra et al., 2009 and Han et al., 2019), whereas CDPKs along with SnRK2s activate the guard cell S-type anion channel (SLAC1) in response to ABA (Geiger et al., 2010; Brandt et al., 2012). Importantly, in the *snrk2.2/snrk2.3/snrk2.6* knockout mutant the Ca^2+^-dependent SLAC1 regulation was impaired (Brandt et al., 2015) indicating interconnection between SnRK2s and calcium signaling pathways. Furthermore, in 2003 Harmon predicted that calcium could somehow regulate SnRK2s (Harmon, 2003); C-terminal parts of SnRK2 kinases are rich in acidic amino acids, and those can be potentially involved in Ca^2+^-binding. Our previous results showed that indeed Ca^2+^ added to SnRK2s slightly affected their kinase activity (Bucholc et al., 2011). However, a much more significant Ca^2+^-dependent effect on their activity arises via their cellular inhibitory partner, SCS, a calcium sensor (Bucholc et al., 2011).

Our present results show that in Arabidopsis, due to alternative transcription start sites, two forms of *AtSCS* are expressed, *AtSCS-A* and *AtSCS-B*. They encode proteins that differ only in the N-terminal region; AtSCS-B is thus a shorter version of AtSCS-A in which the first 110 aa are missing. This region includes the classical EF-hand and the first EF-hand-like motifs. The basic properties of AtSCS-A have been previously described by Bucholc et al. (2011). Now, we analyzed biophysical and biochemical features of AtSCS-B, and performed detailed comparative studies of both proteins in respect of their conformational dynamic with and without calcium, inhibition of the SnRK2 activity and their role in the plant response to water deficit.

The data presented here indicate that only AtSCS-A can play a role as a calcium sensor. The calcium-binding constant of this protein as determined previously by the protein fluorescence titration was 2.5 (±0.9) × 10^5^ M^−1^, corresponding to a dissociation constant of 4 μM (Bucholc et al., 2011). The calcium binding induces significant conformational changes of the protein as revealed by CD spectroscopy and HDX-MS analysis, indicating a possible function of AtSCS-A as a calcium sensor in plant cells. *In vitro* studies demonstrate the SnRK2 inhibition occurs only in the presence of calcium. On the other hand our data showed that even though AtSCS-B binds calcium ions, the binding is two orders of magnitude weaker than that for AtSCS-A and there are no conformational consequences of the calcium binding. These data indicate that for the inhibition of the SnRK2 activity is responsible the common region of AtSCS-A and AtSCS-B. The primary structures of both proteins in this region are identical, than the question appears why AtSCS-A requires Ca^2+^ for the inhibition and AtSCS-B does not. The mechanism of the SnRK2 inhibition by SCS is still an open question and needs further study, but the conformational dynamics of both AtSCSs in the presence and absence of calcium ions shed light on calcium dependence on the SnRK2 inhibition in the case of AtSCS-A. Our HDX-MS results provide extensive analysis of the changes of the conformational dynamic of proteins with classical and non-classical EF-hand motifs upon calcium binding and indicate that N-terminal part of AtSCS-A has a great impact on the C-terminal part.

Our results showed clearly that the domain common to both variants, i.e., the C-terminal fragment of AtSCS-A, is much less stable in the longer form than in the shorter form. HDX-MS and CD analysis indicate that in AtSCS-B this region forms stable structures independent of calcium, with no significant increase in stability in the presence of calcium. In AtSCS-A, the N- and C-terminal segments, though distant in sequence, are structurally coupled, so the C-terminal part has properties distinct from those of AtSCS-B. In AtSCS-A the dynamics of the elements of the C-terminal domain become calcium-sensitive, unlike in AtSCS-B, where calcium-induced changes are negligible. Binding of calcium destabilizes region 321-340 in the C-terminal domain while it reverses the destabilization of the loop of the predicted third EF-hand-like motif (267-278) and the flanking region (278-283). In the presence of calcium ions, the conformational dynamics of this fragment becomes similar to that of the same region in AtSCS-B. Based on these data, we predict that the third EF-hand like motif in AtSCS-A and the motif corresponding to this region in AtSCS-B are probably critical for inhibition of the SnRK2 activity. The conformation of this region required for the kinase inhibition is the one which is present in AtSCS-B (independent of calcium), but which in AtSCS-A requires the presence of calcium. This would explain why AtSCS-A needs calcium ions for the inhibition, whereas AtSCS-B does not.

Moreover, our HDX-MS results show interesting data on the cooperation of Ca^2+^ binding by canonical and non-canonical EF-hand motifs. Upon calcium binding, the region of the predicted calcium-binding loop of the classical EF-hand motif present in AtSCS-A and adjacent regions become much more stable (HDX-MS data), and the overall structure becomes more helical (CD data), Interestingly, according to HDX results, the segment 85-115 (predicted putative EF-hand-like motif with very low identity with the canonical motif) also undergoes significant stabilization in response to Ca^2+^ addition. The changes in deuterium uptake after Ca^2+^ addition in this region were similar to those observed for the canonical motif. It is plausible that the changes observed in the non-canonical motif are not a result of calcium binding in this region, but they are an indirect effect of calcium binding at the canonical site. The vast majority of EF-hand motifs (and almost all fully functional ones) exist in pairs, usually structurally coupled by a short β-sheet (β-scaffold) that maintains a direct contact between metal ion-binding loops (for a review see, Nelson et al., 1998). It is possible that the non-canonical motif cooperates with the canonical motif in calcium ion binding and plays a role of a structural scaffold, influencing the properties of the canonical site. The HDX-MS technique is not able to distinguish between conformational changes resulting from direct ligand binding or allosteric effects triggered by an interaction in a different part of the molecule.

Finally, we analyzed the effect of AtSCS-A and AtSCS-B on plant response to stress using a transgenic approach. We generated transgenic Arabidopsis plants expressing *AtSCS-A-c-myc* or *AtSCS-B-c-myc* in the *scs-1* knockout mutant background. We did not observe any significant differences between the plants of the transgenic lines, the WT plants, and the *scs* knockout mutants grown in optimal conditions. Since SnRK2s are key regulators of the plant response to water deficit (Fujii and Zhu, 2009; Fujita et al., 2009; Nakashima et al., 2009) we analyzed the effect of AtSCS-A and AtSCS-B in the plant response to dehydration. The results showed that both forms of AtSCS are involved in response to this stress; the *scs* mutants were more resistant to dehydration than the WT plants and plants expressing *AtSCS-A-c-myc* or *AtSCS-B-c-myc.* We did not observe significant differences between the transgenic lines expressing one or another form of AtSCS.

The only difference we observed was a less pronounced effect of AtSCS-B than that of AtSCS-A; only in the line B12, in which the expression of AtSCS-B was very high, we observed inhibition of the SnRK2 activity and a meaningful effect on the dehydration response. Importantly, the expression of each of the forms was not able to fully compensate the *scs* mutation, suggesting that both forms are involved in the regulation and their role is not fully overlapping. We obtained similar results using two independent assays, drought tolerance test and water loss in detached rosettes. Based on observed differences in the localization of SnRK2-AtSCS-A and SnRK2-AtSCS-B complexes *in planta* (BiFC assays) we can speculate that both AtSCS forms might regulate phosphorylation of diverse proteins in plant cells. However, it should be noted that the effect of both AtSCSs on the plant response to water withdrawal is similar to but less pronounced than the effects of clade A phosphatases, which are considered as the major negative regulators of the SnRK2s activity.

The analysis of the kinase activity in the transgenic plants showed that both SCSs inhibit the ABA-responsive SnRK2s. Still we cannot exclude that AtSCS-A also inhibits the ABA-non-responsive SnRK2s, since AtSCS-A in contrast to AtSCS-B interacts not only with the ABA-activated SnRK2s but also with kinases of the group 1, which are not activated by ABA. However, the evolution studies performed by Holappa et al. (2017) have shown that SCS proteins appeared in earliest land plants at about the same time as ABA receptors - RCAR/PYR/PYL (RCAR, Regulatory Component of ABA Receptor/PYR1, Pyrabactin Resistance 1/PYL, PYR1-like), PP2Cs (Umezawa et al., 2010; Komatsu et al., 2013; Shinozawa et al, 2019), and ABA-activated SnRK2s (from group 3) that constitute the prototype of the SnRK2 family. The ABA-non-responsive SnRK2s evolved later, in vascular plants. Thus, the ABA-activated SnRK2s, RCAR/PYR/PYLs, PP2Cs, and SCSs seem to consist of ancient regulatory modules of ABA signaling, allowing adaptation to a terrestrial environment.

## CONCLUSIONS

Our studies showed that two isoforms of ASCSs (AtSCS-A and AtSCS-B) are expressed in Arabidopsis. They differ significantly in their expression profiles, calcium binding properties, and conformational dynamics. Both of them inhibit the activity of the ABA-activated SnRK2s and regulate the plant response to water deficit similar to the clade A PP2C phosphatases, although the effect of AtSCSs is not as strong as the one observed in the case of the phosphatases. Moreover, the results provide information on calcium binding properties and conformational dynamics of EF-hand and EF-hand-like motifs present in plant proteins. This extends our knowledge on proteins involved in fine-tuning of the SnRK2 activity in stress signaling in plants, connecting calcium-independent and calcium-dependent pathways.

## MATERIALS AND METHODS

### Plant Materials, Growth Conditions, and Stress Treatments

*Arabidopsis thaliana* lines used in this study include: Col-0 as the wild type background, T-DNA insertion lines; *scs-1* (Salk_051356) and *scs-2* (Salk_104688) previously described (Bucholc et al., 2011), *snrk2.6* (Salk_008068) obtained from the Nottingham Arabidopsis Stock Center, double mutant *snrk2.2/2.3* (GABI-Kat 807G04/Salk_107315) provided by Jian-Kang Zhu, (Purdue University), a quadruple knockout mutant *snrk2.1/2.4/2.5/2.10* (SAIL*_519_C01*/Salk*_080588*/Salk*_075624*/WiscDsLox233E9) described by Maszkowska et. al., 2018 and double mutant *abi1-2/pp2ca-1* (Salk_72009/Salk_28132) obtained from Pedro L. Rodriguez (Instituto de Biologia Molecular y Celular de Plantas). The seeds of *Arabidopsis thaliana* were grown under long day conditions (16-h-light/8-h-dark photoperiod) at 22°C/18°C in soil or in hydroponic culture (Araponics System) as described by Kulik et al., 2012. For expression analysis of *AtSCS-A* and *AtSCS-B* in response to ABA or salt, plants were grown at 21°C/21°C under mid-day conditions (12-h-light/12-h-dark cycle) for 2 weeks in hydroponic culture and seedlings were either mock treated or with 10 μM ABA or 150 mM NaCl, respectively, at specific times. After treatment plants were collected, frozen in liquid nitrogen and stored at -80°C until analyzed. For aseptic cultures, seeds were sterilized in 70% ethanol for 2 minutes then in a water/bleach solution 13:1 (v/v) for 20 minutes. After sterilization, the seeds were extensively washed with sterile water. Seeds were stratified in the dark at 4°C for 3 days. For transient expression experiments, protoplasts were isolated from *Arabidopsis thaliana* T87 cells grown in Gamborg B5 medium as described by (Yamada et al., 2004), six days after subculturing.

### *AtSCS-A* and *AtSCS-B* Expression Analysis in Arabidopsis

AtSCS-A and AtSCS-B expression was analyzed by two-step RT-qPCR. Total RNA was extracted from Arabidopsis seedlings or different organs using a Thermo Scientific GeneJet Plant RNA Purification Mini Kit and treated with Thermo Scientific RapidOut DNA Removal Kit. The efficiency of DNA removal was monitored by PCR with primers for PP2A. First-strand cDNA synthesis was performed using 1.5 μg of total RNA with oligo (dT)_18_ primer and Thermo Scientific RevertAid First Strand cDNA Synthesis Kit according to standard manufacturer’s protocol. Relative expression levels were determined by quantitative PCR in a LightCycler® 480 Roche device, using SYBR GREEN mix (Roche). For each target gene amplification, two gene-specific primers were used (listed in Supplemental Table S1) and all cDNA samples (three replicates) and standards (two replicates) were assayed in a single run. Relative gene expression in each sample was calculated using standard curve method (5-point), normalized using a geometric mean of expression values for two reference genes (*PDF2* and *UBC21*) and scaled to the calibrator sample (Col-0 control).

### Expression of Recombinant Proteins

Expression of recombinant proteins in *Escherichia coli* was performed as previously described (Bucholc et al., 2011). cDNA encoding AtSCS-B or fragment of ABF2 (Gly73 to Gln120) was PCR amplified using appropriate primers (listed in Supplemental Data) and cloned into pGEX-4T-1 vector (GE Healthcare Life Sciences). The PCR reaction was performed using a high-fidelity Pfu DNA polymerase (Stratagene, La Jolla, CA) and verified by sequencing.

### Purification of Recombinant Proteins

All recombinant proteins were purified using glutathione-sepharose beads (GE Healthcare Life Sciences) as previously described by Bucholc et al., 2011. To obtain highly purified AtSCS-A and AtSCS-B proteins the last step of purification was reverse-phase HPLC on an analytical ACT Ace C18 column (for analysis of calcium binding) or Mono Q column using an FPLC system (GE Healthcare Life Sciences). Purity of the proteins was analyzed by SDS-PAGE and electrospray ionization mass spectrometry on a Micromass Q-TOF spectrometer (Micromass, Manchester, Great Britain). GST-ABF2^73-120^ fusion protein after purification using glutathione-sepharose beads (GE Healthcare Life Sciences) was precipitated with chloroform/methanol and resolubized in 10mM Tris-HCL, pH8.8, 0.1% SDS.

### Protein Kinase Activity Assays

The kinase activity assay in solution was performed as described by Bucholc et al., 2011.

In-gel kinase activity assays were performed according to Wang and Zhu, 2016 with some modifications. Proteins were extracted from 14-d-old seedlings grown in hydroponic culture in flasks that were either mock-treated or with 100µM ABA, 350 mM NaCl for 0, 30, 60 min. Plant materials from wild-type (Col-0) and different mutant lines were ground in liquid nitrogen and sonicated three times in the buffer (50 mM HEPES-KOH, pH 7.5, 5 mM EDTA, 5 mM EGTA, 2 mM DTT, 25 mM NaF, 1mM Na_3_VO_4,_ 50 mM ß-glycerophosphate, 10% (v/v) glycerol, 1mM PMSF, 1x protease inhibitor cocktail (Roche) and 1x phosphostop (Roche). Proteins (40 µg/lane) in Laemmli buffer (without boiling) were separated on a 10% SDS-PAGE containing 0.25 mg/mL GST-ABF2^Gly73-Gln120^ or 0.5 mg/mL myelin basic protein (MBP) (Sigma-Aldrich) as a kinase substrate. The gels were run over-night at 30 V and next washed for 3 × 30 min at room temperature (RT) in SDS removal buffer (25 mM Tris-HCl, pH 7.5, 0.5 mM DTT, 5 mM NaF, 0.1 mM Na_3_VO_4_, 0.5 mg/mL BSA, 0.1 % Triton X-100). After that, the gels were incubated in renaturing buffer (25 mM Tris-HCl, pH 7.5, 1mM DTT, 5 mM NaF, 0.1 mM Na_3_VO_4_) for 2 × 30 min at RT, over-night at 4°C and 1 × 30 min at RT. After 30 min of incubation at RT in cold kinase reaction buffer (25 mM Tris-HCl, pH 7.5, 1 mM EGTA, 30 mM MgCl_2_, 2 mM DTT, 0.1 mM Na_3_VO_4,_) the gels were incubated in 10 ml of hot kinase reaction buffer supplemented with 50µCi of [ɤ-^32^P] ATP for 5 min at RT and after addition of 20µM of cold ATP for 90 min at RT. The reaction was stopped with stop washing solution (5% TCA and 1% sodium pyrophosphate). After extensive washing with washing buffer the gels were stained with Coomassie Brilliant Blue R250, dried and exposed to autoradiography.

### Yeast Two Hybrid and Bimolecular Fluorescence Complementation (BiFC) Assays

Yeast two-hybrid analysis was carried out as described previously by Bucholc et al., 2011. The cDNAs encoding SnRK2s were cloned into pGBT9 vector (Clontech) and *AtSCS-A* or *AtSCS-B* were cloned into pGAD424 (Clontech). Primers used for cloning are listed in Supplemental Table S1.

For subcellular localization analysis coding sequences for AtSCS-A and AtSCS-B were introduced into pSAT6-EGFP-N1 or pSAT6-EGFP-C1. For BiFC assays, constructs for *SnRK2s* expression were prepared in pSAT4-nEYFP-C1 and *AtSCS-A* or *AtSCS-B* in pSAT1-cEYFP-N1 or pSAT1-nEYFP-N1, respectively. The pSAT vectors were provided by Prof. T. Tzfira (Tzfira et al., 2005). Primers are listed in Supplemental Table S1. Subcellular localization and BiFC analyses were performed as described by Bucholc et al., 2011. Subcellular localization of the EGFP fusion proteins and EYFP fluorescence after complementation were evaluated using a Nikon C1 confocal system built on TE2000E inverted microscope and equipped with a 60×/1.4 NA Plan-Apochromat oil immersion objective (Nikon Instruments B.V. Europe, Amsterdam, Netherlands). EGFP and EYFP were excited with a Sapphire 488 nm laser (Coherent, Santa Clara, CA, USA) and observed using the 515/530 emission filter. For publication, single optical sections with distinctly visible nucleus and nucleoli were selected to ensure that similar focal planes were compared for all tested variants. Images were processed using Nikon EZ-C1 Free Viewer (version 3.90).

### Calcium Ion Binding Analysis

For calcium binding analysis, the recombinant proteins were purified by reverse-phase HPLC on an analytical ACT Ace C18 column; their purity was estimated by electrospray ionization mass spectrometry. Protein concentration was determined from UV absorption at 280 nm, using the molar extinction coefficients of 10500 M^−1^ cm^−1^ and 9300 M^−1^ cm^−1^ for AtSCS-A and AtSCS-B, respectively, as calculated for one Trp and three Tyr residues (AtSCS-A) or one Trp and two Tyr residues (AtSCS-B) (Mach et al., 1992). The UV absorption spectra were measured on a Cary 3E spectrophotometer (Varian) in thermostated cells of 10 mm path length. 20 mM Tris buffer, pH 7.4, with 100 mM NaCl was used as a solvent for the calcium binding experiments. All measurements were performed at 20°C.

#### Fluorescence Measurements

The protein fluorescence was measured on a CaryEclipse fluorimeter. Spectra were collected with an average time of 2 s for each point and a step size of 0.5 nm from 295 (or 305) to 450 nm. In fluorescence titration experiments small aliquots of concentrated calcium chloride solution were added until the protein fluorescence no longer changed. The measurements were repeated at least three times. All protein solutions used in the fluorescence titration experiments were checked for possible calcium contamination by comparing the protein fluorescence signals in the presence and absence of calcium chelator (100 μM EGTA).

#### Calcium Ion Binding Constant

The measured fluorescence signal (F) is defined as:

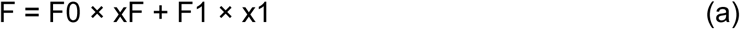

where F0, F1, are fluorescence of the protein without and with the ligand, respectively. The xF and x1, are mole fractions of free protein (PF, no ligand bound) relative to total protein (P0), xF=PF/P0, and ligand bound protein (P1), x1=P1/P0, respectively. The two mole fractions sum to one: xF +x1 =1.

The binding constant of the ligand to the protein in a reaction, P + L ⇄ PL is defined as:

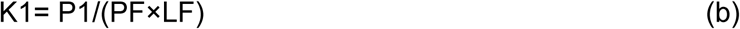

where, P0=P1+PF and L0=LF+P1 are total concentrations of the protein and the ligand, respectively. If F’= F/F0 and f1 = F1/F0, then:

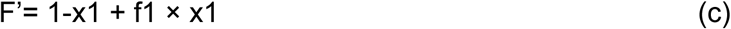

The equation is analogous with that of eq. 8 in (Eftink, 1997) describing single-site ligand binding, where the variables are F’ (from experimental measurements), total ligand concentration, L0, and protein concentration, P0, and the parameters are the ligand binding constant, K1, and relative fluorescence signal, f1. Here, the ligand is Ca^2+^ ion. The parameter values were determined by least-square fitting of theoretical curves to experimental data using the NiceFit program.

### Circular Dichroism Measurements

Circular dichroism (CD) experiments were carried out at 20°C on Jasco J-815 spectropolarimeter with a 10 mm path length cuvette. The protein solutions (close to 2 μM) were prepared in 5 mM Tris buffer, pH 7.4, with 100 mM NaCl. CD spectra were collected twice with an average time of 2 s for each point and a step size of 1 nm from 200 to 270 nm. All spectra were corrected against the buffer. The data were converted to molar residue ellipticity using the relationship [Θ]= θ/(10×n×l×c), where [Θ] is molar residue ellipticity in (degree cm^2^ dmol^−1^), θ is the observed ellipticity in millidegrees, n is the number of aminoacid residues in the protein, l is the path length in cm, and c the protein concentration in M.

The secondary structure content of the proteins was estimated using the CDNN program (CD spectroscopy deconvolution software) (Böhm et al., 1992).

### Hydrogen Deuterium Exchange Coupled to Mass Spectrometry

The HDX-MS was performed as described previously (Sitkiewicz et al., 2013), with minor modifications. Initially, the list of peptides was established by diluting each analyzed protein to 5-10 µM in a non-deuterated buffer (50 mM Tris pH 7.4, 150 mM NaCl). Each sample (50 µL) was acidified by adding 10 μL of “stop” buffer (2M glycine pH 2.5, 150mM NaCl, 4M GuHCl) and then injected into a Waters nanoACQUITY UPLC system equipped with an HDX Manager system (Waters) with the column outlet coupled directly with SYNAPT G2 HDMS mass spectrometer. Rapid online digestion on an immobilized pepsin column (Poroszyme, ABI) was performed at a flow 200 μL/min. Peptides were captured on a trapping BEH C18 1.7-mm, 2.1×5mm Vanguard Pre-Column (Waters) and separated by 1.0×100 mm BEH C18 1.7-mm analytical reversed phase column (Waters) with a gradient of 8-40 % acetonitrile (0.1 % formic acid) in 6 minutes at a flow 40 μL/min. All capillaries, valves, and columns were maintained at 0.5°C inside a HDX cooling chamber, while the pepsin column was kept at 13°C inside the temperature-controlled digestion compartment. Mass spectra were acquired in MSE mode over the m/z range of 50-2000 both with and without ion mobility separation. Spectrometer ion source parameters were as follows: ESI positive mode, capillary voltage 3 kV, sampling cone voltage 35 V, extraction cone voltage 3 V, source temperature 80°C, desolvation temperature 175°C, and desolvation gas flow 800 L/h. Infusion and scanning every 30 seconds of leu-enkephalin (556.277 m/z) was used for continuous lock mass correction. Rigorous washing steps were performed between each injection. The peptides were identified with ProteinLynx Global Server software (Waters) using randomized database and with false discovery rate threshold set to below 4%.

Deuterium labeling was initiated with a 10-fold dilution of the protein sample into a buffer containing 50 mM Tris, pH 7 (uncorrected meter reading), 150 mM NaCl and 5 mM CaCl_2_ or 5 mM EGTA in 99.8% D_2_O (Cambridge Isotope Laboratories, Inc.). After a specified time (10 s, 1 min, 20 min, 60 min) the labeling reaction was quenched by adding “stop” buffer and then the sample was immediately snap-frozen in liquid nitrogen and stored at −80°C until analyzed. Quenched samples were rapidly thawed and injected into a Waters nanoACQUITY LC system equipped with HDX Manager, coupled directly with SYNAPT G2 HDMS mass spectrometer. “Out” controls of back exchange level were performed by incubation of the protein in D_2_O buffer for 48 hours to obtain maximum exchange for each peptide. The experimental maximum is always lower than the maximal theoretical exchange due to a certain degree of back exchange. Each experiment was repeated at least three times and the results represent the mean of all replicates.

#### Data analysis

Peptide list for each protein was created in the DynamX 3.0 software based on PLGS peptide identifications, with following acceptance criteria: minimum intensity threshold - 5000, minimum fragmentation products per amino acids for precursor - 0.3, the maximum mass difference between the measured and theoretical value for parent ions - 10 ppm. Analysis of the isotopic envelopes in DynamX 3.0 software was carried out using the following parameters: retention time deviation ± 18 s, m/z deviation ± 15 ppm, drift time deviation ± 2 time bins and centroids of the mass envelopes were obtained. The values reflecting the experimental mass of each peptide in all possible states, replicates, time points and charge states were exported from the DynamX 3.0 and further data analysis was carried out using in house scripts written in R language (http://www.R-project.org). Fraction exchanged (D) was calculated with the following formula:

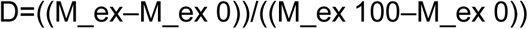

where: (Mex0) and (Mex100) indicate the average peptide mass measured in the unlabeled sample and mass from “out” control, respectively. Error bars for fraction exchanged represent standard deviations calculated from at least three independent experiments. The difference in the fraction exchanged (Δ fraction exchanged) was calculated by subtracting the fraction-exchanged values for peptides in the selected state from the values for the same peptides in the control state. The error bars were calculated as the square root of the sum of the variances from compared states. Student’s t-test for two independent samples with unequal variances and unequal sample sizes (also known as Welsh t-test) was carried out to evaluate differences in fraction exchanged between the same peptides in two different states.

### Construction and selection of transgenic Arabidopsis thaliana plants

To generate the transgenic lines in *scs-1* background coding sequences for AtSCS-A and AtSCS-B were cloned into pENTR-®-D/TOPO™ vector (Thermo Fisher) and next, the cDNAs were recombined by Gateway LR reaction into pGWB617 destination vector (Nakamura et al.,2010). The pGWB617-AtSCS-A and pGWB617-AtSCS-B constructs were transformed via *Agrobacterium tumefaciens* strain GV3101 into mutant background *scs-11* by the floral dip method as previously described by Clough & Bent, 1998 and Zhang et al., 2006. The selection of the transgenic lines was performed by spraying soil-grown seedlings with 0.033% BASTA solution supplemented with 0.01% Silwet L-77 at 7 and 9-day after germination. Recombinant proteins were confirmed by western blot analysis using a c-Myc monoclonal antibody (9E10, Santa Cruz) according to the protocol recommended by the manufacturer.

### Rosette water loss measurement

The Arabidopsis plants of the appropriate genotype were grown for 6 weeks under short day conditions (8 h light at 22°C / 16 h dark at 20°C) in a CLF PlantClimatics chamber incubator and watered copiously one day before harvest. The Cut Rosette Water Loss (CRWL) was determined as described previously by Bouchabke et al. (2008) with minor modifications. Freshly cut rosettes were weighed immediately, incubated in windless conditions under constant temperature (22-24°C) and weighed five times hourly. After overnight drying at 70°C to a constant mass, the rosettes were weighed for dry mass, and water loss was calculated.

### Drought tolerance test

The Arabidopsis plants were grown in pots (approximately 50 plants per pot) for 17 days under long-day conditions (16 h light at 22°C / 8 h dark at 20°C) and for an additional 2 weeks without watering. After that time, the plants were watered. Pictures were taken before re-watering and on the second day of re-watering.

### Modeling of AtSCS-A/B Structure

Since no structures of AtSCS variants have been available in the PDB, they were modeled by homology using algorithms implemented in Yasara (Krieger et al., 2002; Krieger et al., 2009). In every round of modeling 12 protein structures sequentially close to the target were used as initial templates. For each template up to 5 alternative alignments with the target sequence were used, and up to 50 different conformations were tested for each loop being modified. The resulting models were individually scored according to their structural quality (dihedral distribution, backbone and side-chain packing), and those with the highest scores were further used to form a hybrid model, which was built using the best fragments (e.g. loops) identified among the single-template models. However, due to limited coverage with sequences of proteins with known structure, AtSCS-A has to be modeled as the two separate domains, relative orientation of which remains unknown. In accordance with HDX-MS data, the N-terminal one covered residues 26-175, while the C-terminal corresponded to residues 211-330.

## ACKNOWLEDGMENTS

We thank dr. Olga Sztatelman (IBB PAN, Warsaw, Poland) for providing pGBT9-SnRK2.3, pGBT9-SnRK2.7, pGBT9-SnRK2.9 plasmids and *snrk2.1/4/5/10* knockout mutant. We are also grateful to Professor Zhu-J-K and Pedro l. Rodriguez for sharing with us the *snrk2.2/2.3* and *abi1-2pp2ca-1* knockout mutants, respectively, and all members of our laboratory for stimulating discussions. We also thank Nottingham Arabidopsis Stock Center for providing the *snrk2.6* mutant.

## Supplemental Data

Supplemental Table S1. List of primers.

Supplemental Figure S1.

AtSCS-A *and* AtSCS-B *transcript levels in different organs*.

Different presentation of data showed in Fig.1.

Supplemental Figure S2

*AtSCS-A-c-myc and AtSCS-B-c-myc protein level in homozygous 35S::AtSCS-A-c-myc and 35S::AtSCS-B-c-myc transgenic plants.* The level of AtSCS-A-c-myc and AtSCS-B-c-myc was analyzed by Western blotting using anti-c-myc antibodies, as described in Materials and Methods.

Supplemental Figure S3.

*Analysis of expression of* Rab18 *and* RD29B *in Arabidopsis plants with different level of AtSCS-A and AtSCS-B.* The relative transcript level of *RAB18* and *RD29B* in RNA extracted from 2-week-old seedlings that were either mock or 10 µM ABA-treated for 3 h was analyzed by quantitative RT-PCR

Supplemental Figure S4.

*The stoichiometry of the binding of calcium ions to AtSCS-B.* The index n equal to 1 was calculated based on the general equation describing the association of the ligand to the protein used for the analysis of the fluorescence measurements. The squares and the rings describe the data from two independent experiments.

Supplemental Figure S5.

*Differences in the Hydrogen/Deuterium (H/D) exchange pattern in AtSCS-A and AtSCS-B upon Ca^2+^ binding.* Differential plot of H/D exchange presented in Fig. 8. Δ Fraction Exchanged was calculated by subtracting Fraction Exchanged values in peptides of AtSCS-A (left panels) and AtSCS-B (right panels) in the presence and absence of calcium. Peptides with p-values below 0.01 in the t-test are labeled in bright purple. Dark grey – position of calcium binding loops of the canonical EF-hand motif, light grey – positions of calcium binding loops of potential non-canonical EF-hand motifs.

Supplemental Figure S6.

*Differences in the Hydrogen/Deuterium (H/D) exchange pattern between AtSCS-A and AtSCS-B in common regions in the presence and absence of calcium ions.* Δ Fraction Exchanged was calculated by subtracting Fraction Exchanged values in the same peptides of AtSCS-A and AtSCS-B in the presence and absence of calcium, respectively (samples are the same as presented in Fig. 8). Peptides with p-values below 0.01 in the t-test are labeled in bright purple. Light grey – positions of calcium binding loops of potential non-canonical EF-hand motifs.

Supplemental Figure S7.

*Kinetics of H/D exchange in AtSCS-A and AtSCS-B in the presence or absence of Ca^2+^.* Kinetics of H/D exchange in the presence of Ca^2+^ (magenta lines for AtSCS-A, green lines for AtSCS-B) and in the absence of Ca^2+^ (blue lines for AtSCS-A, black lines for AtSCS-B) for peptides resulting from AtSCS-A fragments starting from residue 2 to residue 81 (A), from AtSCS-A and AtSCS-B fragments from residues 85 to 160 (B), from AtSCS-A and AtSCS-B fragments from 148 to 255 (C), and from AtSCS-A and AtSCS-B fragments from 245 to 375 (D). H/D exchange was analyzed at four time points plotted on a logarithmic scale. Time in minutes (10 sec, 1 min, 20 min, 60 min) is shown at the top of the graph. Error bars represent standard deviations calculated from at least three independent experiments.

Supplemental Figure S8.

*Model of N-terminal domain of AtSCS-A (region 16-119) (A) and C-terminal domain of AtSCS-A and AtSCS-B (region 241-339) (B) with overlaid H/D exchange results.* Regions with relatively high H/D exchange (exchange fraction higher than 0.5 after 1 min incubation with D_2_O in the presence of Ca^2+^) are marked with yellow, regions with retarded H/D exchange (exchange fraction lower than 0.5 after 1 min incubation with D_2_O in the presence of Ca^2+^) are marked with purple. For AtSCS-A kinetics of H/D exchange is presented in the presence (red) and absence of Ca^2+^ (blue) (A) and (B), for AtSCS-B in the presence of Ca^2+^ (green) and absence of Ca^2+^ (black) (B).

